# A complex reach direction rule that delays reaction time causes alternating excitation and inhibition in express muscle responses and corticospinal excitability

**DOI:** 10.1101/2022.12.13.520353

**Authors:** Rechu Divakar, Gerald E. Loeb, Brian D. Corneil, Guy Wallis, Timothy J. Carroll

## Abstract

The dynamics of muscle activation during fast visually guided reaching are suggestive of two neural control signals; an early signal that acts at “express” latencies in response to the visual stimulus, and a longer latency signal that executes a strategic reach plan. Here we developed a task designed to temporally isolate the express visuomotor response from the longer latency muscle response, and to characterize the time course of corticospinal excitability changes in the express response time window when the late voluntary response is delayed. We tested this by measuring electromyograms (EMG) and changes in Motor Evoked Potential (MEP) amplitudes following Transcranial Magnetic Stimulation (TMS) of the motor cortex, as participants reached either towards or away from visual targets. Crucially, the information about the task rule was provided by the luminance of the target itself, and so was unknown to the subject until the instant of target presentation. This feature delayed reaction times, likely because additional (presumably cortical) processing was required to interpret and apply the rule before formation of a goal directed reach plan. The earliest EMG responses to target presentation occurred with a 70-105 ms time window, and were oriented to bring the hand toward the location of the target. However, there was also a slightly later response that was also time-locked to target appearance in a 105-140 ms time window. This second response was “reciprocal” to the first, such that it was oriented to take the hand in the direction opposite from the target. In some participants, additional oscillating cycles were apparent after the first two target-related responses. These multiphasic express visuomotor responses were nearly identical in both pro- and anti-reach conditions. These muscle activity responses were generally reflected in the temporal pattern of corticospinal excitability modulations in experiment two. Indeed, the MEP and background EMG responses showed an alternating pattern similar to that in experiment one, although the effect was clearer in the anti-reach than the pro-reach condition. Overall, the data show that the express and voluntary responses are indeed distinct neural control signals, which supports the hypothesis that at least two separate neural pathways (one slow and one fast) contribute to the control of visually guided reaching. The properties of the fast pathway are consistent with a tecto-reticulospinal pathway, while those of the slow pathway are consistent with a transcortical loop.

## INTRODUCTION

Reaching to a visual stimulus involves activating agonist and inhibiting antagonist muscles relevant for the reach. Electromyographic (EMG) activity of the muscles involved typically shows patterns of activation or inhibition that are distributed around the initiation of arm movement, with the onset of EMG correlating with the onset of the arm movement. However, around 80 milliseconds after the onset of a visual stimuli, muscles involved in the reach can be activated or inhibited in a manner that is not correlated with the onset time of the arm movement but is instead time-locked to onset of the visual stimulus. This “express visuomotor response” may be driven by subcortical pathways which include the superior colliculus and reticular formation (Corneil and Munoz, 2014). The “late muscle response” that is associated with the movement onset and that presumably triggers the arm movement is widely thought to be driven by transcortical pathways (Kalaska, 2009). Thus, the possibility of two neural pathways exists – one that acts on the muscle at express latencies in response to the visual stimulus, and another that acts on the muscle at longer latencies in response to a strategic reach plan.

Strong express visuomotor responses (EVRs) are typically elicited when the visual target is temporally and spatially predictable (Contemori et al., 2020). Thus, most studies investigating the EVR used paradigms with partially predictable targets, which inadvertently causes shorter reaction times. Consequently, EVRs within the 70-120 ms window often merge with the late muscle responses that can start anywhere from 120-250 ms post stimulus. This immediate or overlapping transition from one response to the other limits our ability to study the origins of these putatively distinct responses in isolation. The aim of this study, therefore, was to 1) devise a way to temporally isolate the express visuomotor response from the late muscle response, and 2) characterize the time course of corticospinal excitability changes in the express response time window when the late voluntary response is delayed. If we can temporally isolate the EVR and associated excitability changes from the late response, it would also add to the evidence for the existence of two separate neural pathways that work at different timescales.

To this end, we adopted the emerging target reaching paradigm developed by Kozak et al. (2020) and modified it to increase mechanical reaction time. Participants executed fast reaches either in the direction of (pro-reach) or in the direction opposite (anti-reach) of an abruptly appearing stimulus. Crucially, the information about whether to execute a pro-or anti-reach was encoded in the luminance of the target. Thus, the task rule was unknown to the subject until the instant of target presentation. Withholding information about the rule that defines the task-goal until the appearance of the stimulus was expected to slow down reaction times, as additional (presumably cortical) processing would be required to interpret and apply the rule (Fischer and Weber, 1992; Carey et al., 1996; Klein and Foerster, 2001; Heath et al., 2009). Increasing task complexity has been shown to reduce the amplitude of express visuomotor responses (Gu et al 2018), therefore, it was important to test whether this modified paradigm could elicit express responses despite the complexity.

The first experiment tested whether a cognitively demanding motor task can produce express muscle responses that are temporally isolated from the voluntary muscle response. The second experiment measured corticospinal excitability changes within 60-120 ms of target onset, when later muscle response signals associated with the onset of the movement were delayed. We tracked the changes in amplitude of motor evoked potentials (MEPs) in right-side upper limb muscles by stimulating the left motor cortex at various time points after the onset of the visual stimulus. As the modified paradigm was expected to delay the onset of the movement, we expected to see delayed movement-related excitability changes. The purpose was to reveal the timing and duration of excitability changes associated with the express response without contamination from movement-related inputs. A clear dissociation between excitability changes for the two muscle response types would further support the notion that the express and late voluntary responses are produced by different neural substrates.

## METHODS

### Participants

Twenty-one participants (8 males, 13 females; mean age: 28.7, SD: 5.6; 18 right-handed, 3 left-handed) took part in the first experiment and twenty-one participants (7 males, 14 females; mean age: 25.1, SD: 6.7; all right-handed) took part in the second experiment. Three participants who took part in experiment one also participated in experiment two. Participants provided their informed consent after they read a participant information document and received a verbal explanation of the procedures. For the second experiment, subjects also filled out a health screening questionnaire to make sure they were not contraindicated for TMS stimulation. All participants were free to withdraw from the experiment at any time. The experimental procedures conformed to the Declaration of Helsinki, and the University of Queensland Medical Research Ethics Committee (Brisbane, Australia) approved the procedures.

### Experimental design

The general setup used in the study (**Figure 1A**) has been described previously (Divakar et al., 2022). Briefly, in both experiments 1 and 2, we used an emerging target paradigm (**Figure 1B**) that has been shown to promote short latency muscle responses (Kozak et al., 2019; Contemori et al., 2020). After a 500 ms unbroken fixation was detected, a visual target moved down an inverted y-shaped track and gradually disappeared behind a rectangular barrier. It then briefly reappeared beneath the barrier (as a complete circle) on either the left or right side of fixation. On a given trial, participants had to reach quickly either toward (pro-reach) or away (anti-reach) from the target. All movements were executed by either shoulder extension (right reaches) or shoulder flexions (left reaches) in the transverse plane.

**Figure 1.**
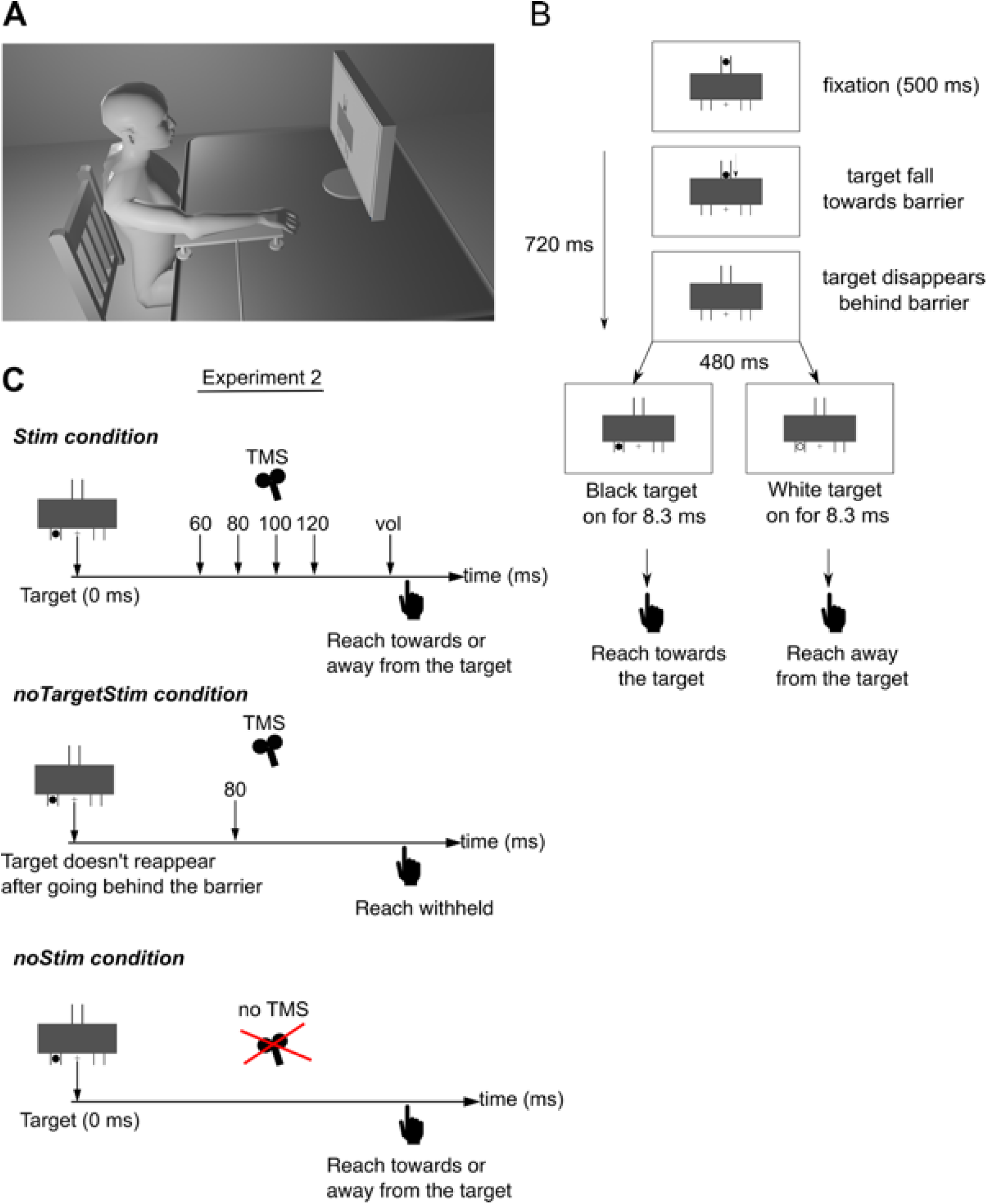
***A*:** Experiment setup. Participants sat facing a monitor, their right arm resting on an air-sled. A custom-made pulley mechanism attached to the air-sled applied a load force of ∼5N (towards right) to increase the baseline activity of the pectoralis major muscle. In the first experiment, EMG data were recorded from the pectoralis major and posterior deltoid and in the second experiment data was recorded from pectoralis major (PEC) of the right arm. An accelerometer attached to the right index finger recorded the onset of the arm movement. ***B*:** In both experiments, we used an emerging target paradigm, in which a visual target moves down an inverted y-shaped track at a constant speed, and gradually disappears behind a rectangular barrier. It then briefly reappears beneath the barrier on either the left or right side. On a given block, participants were instructed to either quickly reach for (pro-reach) or away (anti-reach) from the target depending on the luminance of the target. In the first experiment, participants reached towards the target if it was black and reached away if it was white. In the second experiment, the link between luminance and reach-type (pro or anti) was counterbalanced across participants. ***C*:** In the second experiment, a TMS pulse was delivered after the target onset (Stim condition) on a subset of trials. TMS was delivered to the left motor cortex at either 60, 80, 100, 120 or at a “voluntary” latency (30 millisecond prior to the median onset latency of the voluntary muscle activity). Brain stimulation was also delivered on some trials when target did not reappear beneath the barrier (noTargetStim condition), and in the remaining trials no stimulation was delivered after the target appeared (noStim condition).

The first experiment tested whether short-latency muscle responses could be temporally separated from the long-latency muscle responses. We used an anti-reach paradigm in which the instruction regarding whether to execute a pro-reach (towards the target) or an anti-reach (away from the target) was embedded in the luminance of the visual target. If the target was black, participants were instructed to reach for it and if the colour was white, participants were instructed to reach away from it (**Figure 1B**). Similar paradigms where task instruction is embedded in the target have been shown to increase the reaction time of the participants during reaching (Westendorff and Gail, 2011; Gu et al., 2016) and saccade behaviour (Fischer and Weber, 1992; Everling et al., 1998; Pierce and McDowell, 2016). Participants performed 6 blocks of 60 trials each (360 trials total), in which pro- and anti-reach trials were randomly intermixed. Thus, there were 90 trials each for the 4 conditions – 2 targets (left and right) by 2 reach-types (pro- and anti-reach).

The second experiment tested whether visual target information influences early corticospinal excitability when movement-related outputs have been delayed. We used a similar paradigm to experiment one, with the addition of transcranial magnetic stimulation (TMS). In this experiment, the target luminance representing the pro-or anti-reach rules was counter-balanced across participants. Single magnetic pulses were delivered to the motor cortical areas that generated consistent motor evoked potentials (MEPs) from the clavicular head of the pectoralis major muscle (PEC). For pro- and anti-reaches, single magnetic pulses could be delivered at 6 potential latencies after the onset of the visual target. Five of these latencies were fixed and common for all subjects at 60, 80, 100, 120 and 140 ms (**Figure 1C**). Only half the participants (N = 11) were stimulated at the 140 ms latency. The last stimulation latency, termed the *voluntary* latency, was calculated by subtracting 30 ms from the median onset latency for pro- and anti-reach EMGs. Since the muscle onset times varied between participants, the *voluntary* latencies for pro- and anti-reach trials were unique for each participant. To calculate the *voluntary* latencies participants performed two blocks of (40 trials each) the paradigm before the start of the TMS blocks. Trials in which a TMS pulse was delivered were called ‘*Stim*’ trials (**Figure 1C**).

Two other types of trials were also intermixed along with the *Stim* trials. One type was called ‘*noStim*’, in which no magnetic stimulation was delivered after the onset of the visual target (**Figure 1C**). The second type was called ‘*noTargetStim*’, in which stimulation was delivered without a target. The target would move down the track and disappear behind the barrier, but it would not reappear. The TMS pulse was delivered 720 ms after target started dropping towards the barrier, or 80 ms after the expected “target onset” had the target appeared beneath the barrier (**Figure 1C**). In this way, we could investigate how the TMS pulse influenced the corticospinal tract when the participant was prepared to move but received no target information. Subjects were unaware of when the *noTargetStim* trials would occur and were instructed to withhold the reach if no target appeared under the barrier. The purpose of these trials was to understand the contribution of visual target information to corticospinal excitability, as opposed to potential excitability changes caused by target expectation and motor preparation. To summarize, the *Stim* trials had 24 conditions: 6 latencies (60, 80, 100, 120, 140, vol) by 2 targets (left and right target) by 2 tasks (pro and anti-reach), and the *noStim* trials had 4 conditions: 2 targets (left and right target) by 2 tasks (pro and anti-reach). For each participant, there were 28 trials for each *Stim* condition, 64 trials for each *noStim* condition, and 30 trials in the *noTargetStim* condition. Trials were split as 14 blocks of 66 trials each. On each block, participants encountered 48 *Stim* trials, 2 *noTargetStim* trials, and 16 *noStim* trials, and these trials were randomly intermixed.

### Brain Stimulation

Transcranial Magnetic Stimulation (TMS) was delivered using a figure of eight coil (wing diameter of 8 cm) connected to a MagStim 200^2^ stimulator (The MagStim Company, Carmartheshire, Wales, UK). To find the optimal location on the scalp that produced consistent motor evoked potentials (MEPs) from the pectoralis major muscle we first measured the participant’s head to locate the Cz electrode location (EEG 10-20 system). With the center of the TMS coil starting from a spot 2 cm anterior and 3 cm lateral to the Cz site, single pulses were delivered to the scalp. We adjusted the coil position and stimulator output intensity until consistent MEPs could be induced from the pectoralis major muscle. The coil was held tangential to the scalp with the handle pointing backwards and oriented approximately 45° with respect to the sagittal plane. Note that the pectoralis muscle was loaded using a pulley, which increased the baseline activity of the muscle and lowered the stimulation intensity required to induce an MEP (Di Lazzaro et al., 1998). An Arduino Uno development board in conjunction with a photodiode was used to ensure precise timing of stimulation. The photodiode sent a pulse to the Arduino board when it detected a visual target. The Arduino board then triggered the TMS or TES device at the stimulation latency provided to it by the experiment computer at the start of every trial.

### Data Recording

In the first experiment, surface EMG were recorded from the clavicular head of pectoralis major muscle (PEC) and the posterior head of the deltoid (PD) using a double-differential surface electrodes (Bagnoli-8 system; Delsys Inc.,Boston, MA). The EMG signals were amplified (x1000) and band-pass filtered between 20 Hz and 450 Hz using the native Delsys Bagnoli-8 Main Amplifier Unit. In the second experiment, EMG were recorded from the clavicular head of the pectoralis major muscle (PEC) of the right arm using 24 mm disposable Ag-AgCl electrodes. The EMG signals were amplified (x1000) and band-pass filtered between 30 Hz and 10000 Hz using Grass P511 isolated amplifier. We could not use the Delsys electrodes in the second experiment because the transcranial magnetic stimulation causes large artifacts in the EMG data if recorded using a low pass filter of 450Hz as fixed in the Delsys system. A three-axis accelerometer (Dytran Instrument, Chatsworth, CA) placed on the right index finger measured the acceleration of the arm; these data were used to detect the onset and direction of the arm movement. EMG and accelerometer data sampled at 5000 Hz were digitized and stored on a computer using a 16-bit analog-digital converter (USB-6343-BNC DAQ device, National Instruments, Austin, TX). Eye tracking data were sampled at 1000 Hz with an EyeLink 1000 system (SR Research Ltd., Ontario, Canada). Trials were only initiated upon a stable period of eye fixation (500 ms), and trials in which fixation was broken before target appearance were aborted and redone.

### Data Analysis

Reaction time was calculated by first converting the acceleration data to velocity using the cumulative integral function (cumtrapz) in MATLAB. For a given reach, reaction time (RT) was determined by first identifying the point of peak velocity and then going backward in time to the nearest point when velocity reached 5 % of peak. Only the first peak which corresponded to the initial arm movement was considered for picking the peak velocity. Reach direction was identified by checking the relative sign of the velocity curve (positive indicates a left reach and negative indicates right reach) at peak velocity. To minimize the influence of the movement related EMG in the express response time window, we removed trials with reaction times less than 140 ms from both experiments.

MEP amplitude was calculated by first extracting a 150 ms window (−50 ms to 100 ms with respect to the stimulation onset) around the stimulation. The amplitude of an MEP was then calculated on a trial-by-trial basis by taking the mean of the rectified MEP waveform within the defined MEP window. The MEP window for each muscle varied slightly across participants. For every participant, the start and end latency of the MEP window in a muscle was assigned by visual inspection of the average MEP trace. To compare across participants, we represented the MEP amplitudes of each participant as a percentage their average MEP size in the *noTargetStim* trials.

To calculate the underlying muscle activity or background EMG, in the absence of TMS, we looked at the muscle activity in the no stimulation trials (*noStim* trials) that were intermixed with the stimulation trials. As in detailed in Divakar et al. (2022), we measured the average of the rectified EMG in the *noStim* trials from the same time window that was used to calculate the MEP in the stimulation trials. This time window was called the ‘measurement window’. To compare across participants, the background EMG amplitude was expressed as a percentage of the muscle activity in the *noTarget* trials in a 25 ms window before stimulation onset.

Statistical analysis was performed in GraphPad Prism (Version 9.4.0, GraphPad Software, San Diego, California US). We predominantly used the repeated measures ANOVA to analyze the independent variables. The Greenhouse-Geisser correction was applied whenever Mauchly’s test of sphericity was violated. We inspected the QQ plots to check whether data followed a normal distribution. All distributions were approximately normal and parametric statistics were used throughout. For analyzing the MEP and background EMG data in the second experiment, we used a linear mixed-effects model based on the maximum likelihood method. This was necessary as only half of the subjects were stimulated at 140 ms, and this model helped us to account for the missing values. When there was a significant effect, we used Holm-Šídák tests for post-hoc comparison. For all tests, statistical significance was set at P < 0.05.

## RESULTS

### Experiment 1

The primary objective of experiment 1 was to delay the mechanical reaction time and isolate the express visuomotor response. To attain this objective, we withheld the information about the reach-type (pro-or anti-reach) from the participant until the instant that the target was presented. The luminance of the target (black or white) conveyed this information, which means that participants had to wait until the target appeared and then decide whether to reach towards or away from it depending on the luminance. This was expected to reduce task predictability and delay the reaction time.

Task performance in experiment one shows that the modified paradigm was successful in delaying the mechanical reaction time and the associated voluntary EMG. We saw a median reaction time of ∼330 ms [range: 238-428] ms in this experiment. This was ∼90 ms slower than reaching paradigms where pro- and anti-trials were grouped into separate blocks such that participants knew the reach rule in advance of target presentation (Contemori et al., 2020). Furthermore, participants generally reacted faster in pro-reach (median: ∼311 ms, range: 225– 454 ms) than anti-reach (median: ∼358 ms, range: 245–436 ms) trials (Two-way RM ANOVA, *F*_1,20_ = 45.4, *P* < 0.001, **Figure 2A**). While there was no significant main effect of target direction (*F*_1,20_ = 1.7, *P* = 0.207), there was a significant target direction by reach-type interaction (*F*_1,20_ = 8.8, *P* = 0.008). However, post-hoc comparisons showed that within a given reach-type, reaction time did not differ significantly between left and right targets (**Figure 2A**). The number of reach errors in which participants did not reach according to task instruction was neither significantly influenced by the reach type (*t*_20_ = 0.3, *P* = 0.802) or target direction (*t*_20_ = 0.5, *P* = 0.658). On average participants made 16 (range: 3 to 42) reach errors out of 360 trials.

**Figure 2.**
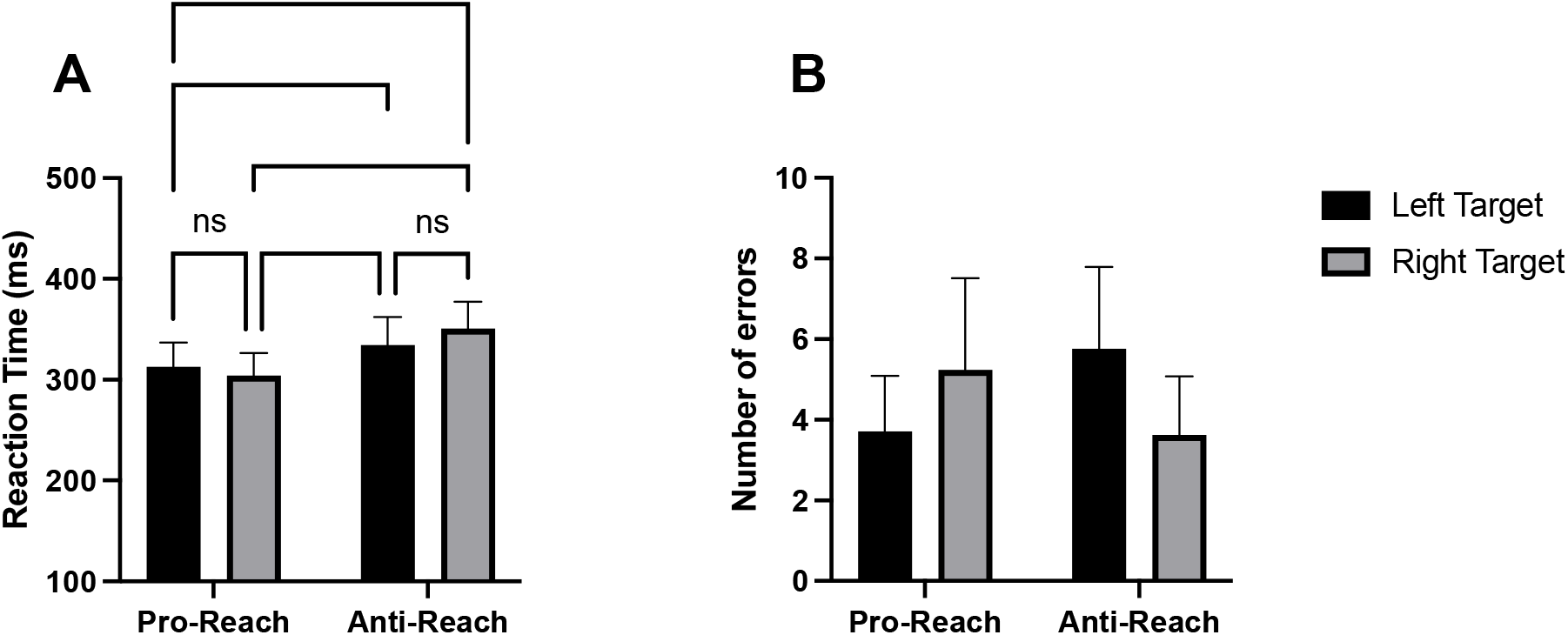
Reaction time across all subjects. ***A*:** Average reaction time in each condition. ***B*:** Average number of reach errors in each condition. * P < .05, ** P < .01, *** P < .001

Previous work showed that the prevalence of express visuomotor responses is strongly dependent on the predictability of the visual stimulus, with the rate dropping as the task becomes less predictable (Contemori et al., 2020). Since our paradigm offered minimal predictability, we checked whether overt express visuomotor responses were produced in this paradigm. **Figure 3** shows the EMG activity in the pectoralis muscle of an exemplar participant, where we see muscle behaviour that is strikingly different from the stereotypical express visuomotor responses reported previously in the 70-120 ms time window. Typical express visuomotor responses in the pectoralis major muscle are characterised by muscle activation when the left target is presented and muscle inhibition when the right target is presented (Gu et al., 2016). In line with typical behaviour, our exemplar participant showed muscle activation in the 70-105 ms time window (shown by black arrow above the raster) for the left target in both pro- and anti-reach condition (**Figure 3A, B, C**), but contrary to typical behaviour, this participant did not show muscle inhibition in this window when the right target was presented. Instead, for right targets, the muscle responded with strong activation in the 105-140 ms time window (red arrow above the raster) in both pro- and anti-reach condition (**Figure 3D, E**). Left targets generated muscle inhibitions (red arrow above the raster) in this time window (**Figure 3A, B**). Thus, the direction of the muscle activity in the 105-140 ms time window appear to be “reciprocal”, or in the direction opposite from the initial target-oriented response in the 70-105 ms time window.

**Figure 3.**
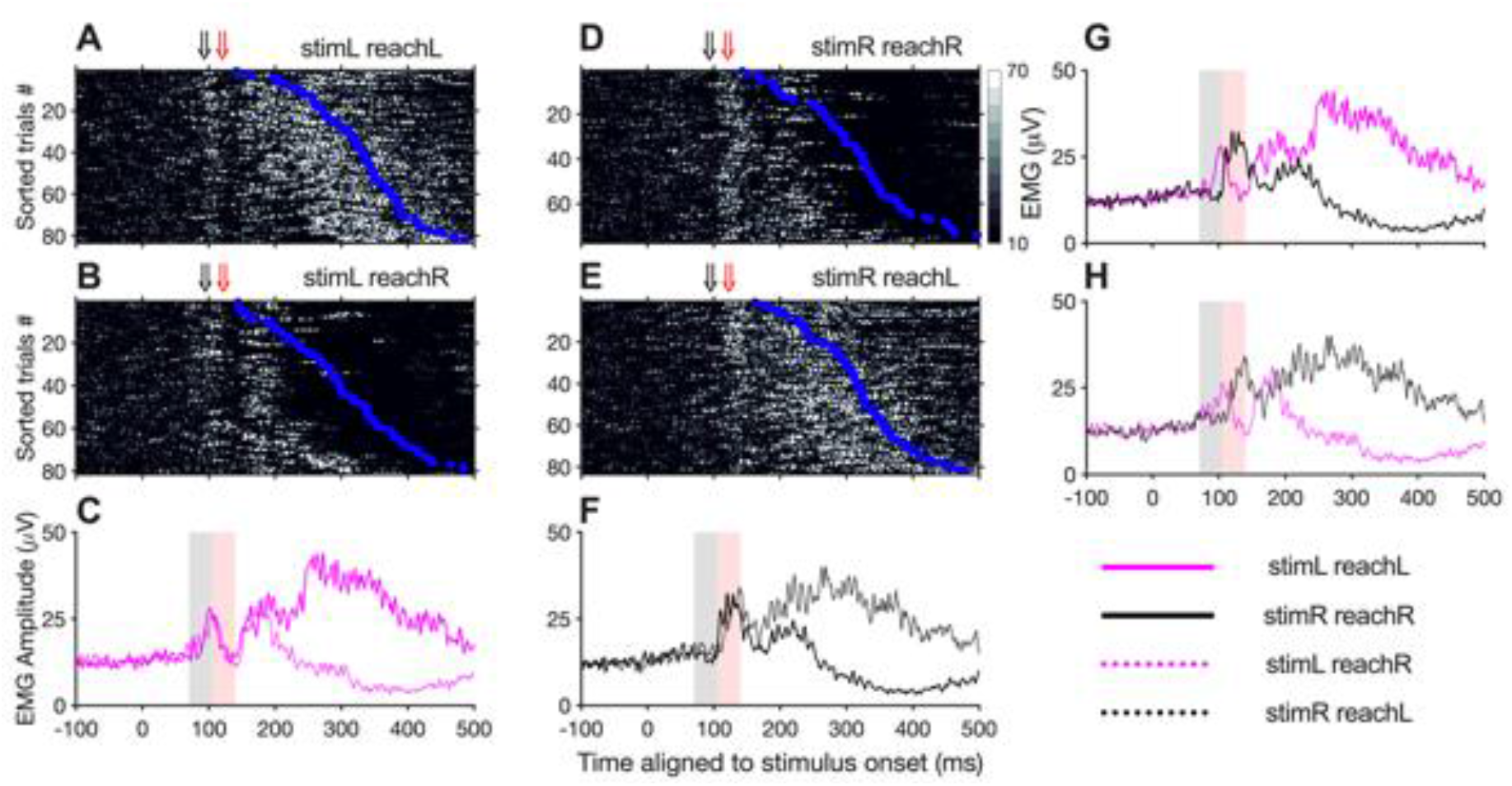
Rectified surface EMG recordings from pectoralis major muscle of an exemplar subject. EMG activity for correct pro-reach trials for left (**A**) and right targets (**D**). Each row of the raster plot represents activity within a single trial, trials are sorted based on the reach reaction time (blue circles) and aligned to the onset of the visual stimulus. EMG activity for correct anti-reach trials for left (**B**) and right targets (**E**). Black arrows above the raster represent the 70-105 ms target-oriented express response window and red arrows represent the 105-140 ms “reciprocal” response window. Average EMG activity of left (**C**) and right target (**F**) directions in pro-(solid line) and anti-reach (dotted line) trials. Average EMG activity of pro-(**G**) and anti-(**H**) trials for left (magenta line) and right (black line) targets. In the average plots, black shaded regions (70-105 ms) show the express response window and red shaded regions show the reciprocal response window (105-140 ms).

Raster plots from five additional subjects (**Figure 4**), show that this reciprocal activity was consistent across participants. In fact, the later reciprocal muscle response was often stronger and more consistent than the initial target-oriented response. Some subjects showed a clear target-oriented response (**Figure 4A, C, D**), whereas this first response was almost absent in others (**Figure 4B, E**). Regardless of the presence or strength of the initial response (70-105 ms), the reciprocal muscle response (105-140 ms) was consistently present. Surprisingly, the reciprocal muscle response was stronger for the right targets compared to the left targets (compare *stimR*, columns 2&4 vs *stimL*, columns 1&3 in **Figure 4**). When the right target was presented the pectoralis muscle was strongly activated regardless of whether it was agonist or antagonist for a particular reach (*stimR* - column 2&4). Such biased responses for right targets have been previously reported (Wood et al., 2015).

**Figure 4.**
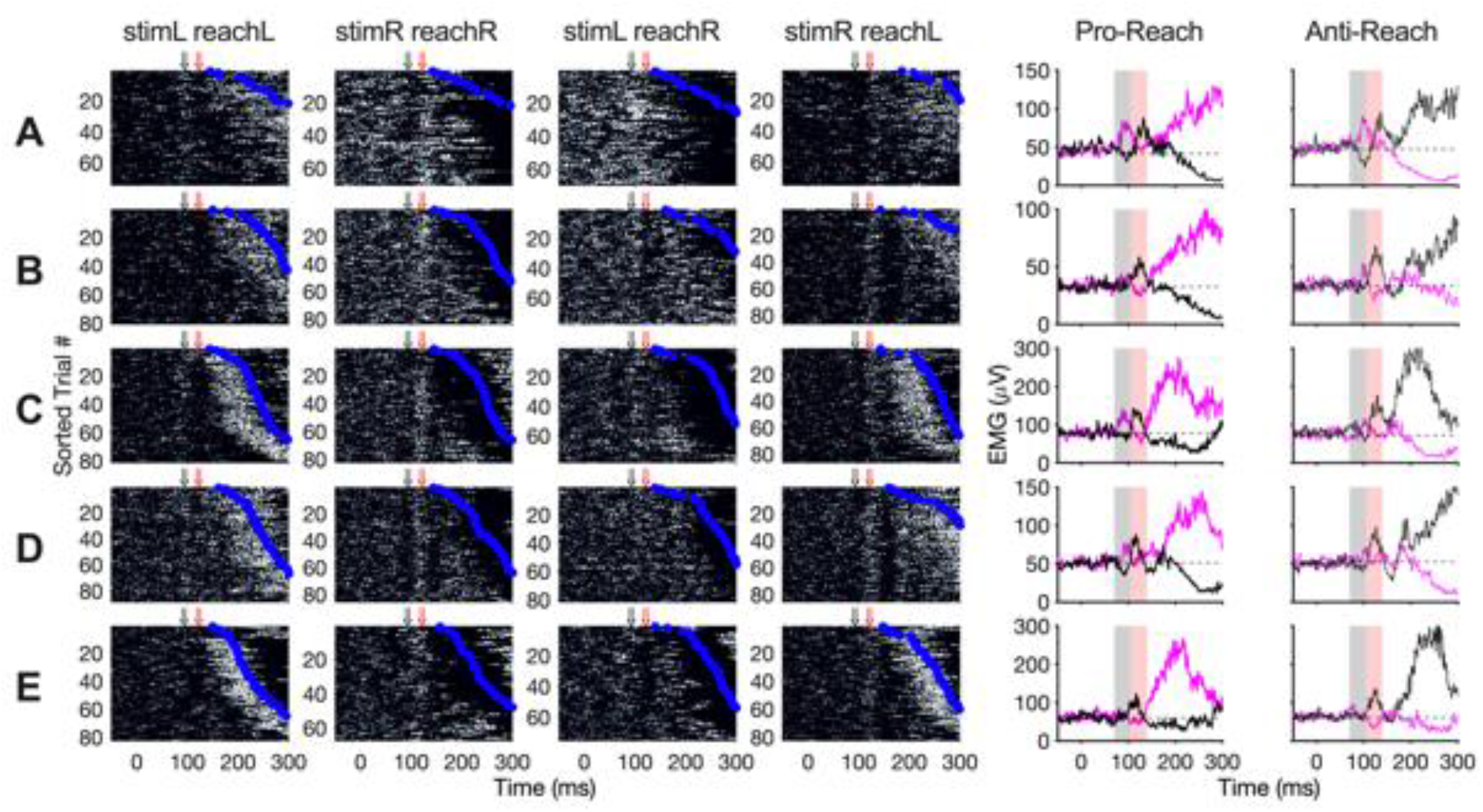
EMG recording from five participants (each row represent data from one participant) showing varying degrees of express target-oriented response in the 70-105 ms time window (shaded grey in average plots) and reciprocal response in the 105-140 ms time window (shaded red in average plots). Plots follow the same convention as in Figure 3.

The group average EMG from the pectoralis muscle shows that when the voluntary movement was delayed the muscle activity in the early phases of reaching (prior to 140 ms) was similar in both pro-reach and anti-reach conditions (**Figure 5A, B**). A two-way RM-ANOVA showed that EMG activity strongly depended on the interaction effect (*F*_3,60_ = 43.70, *P* < 0.001) between time window (70-105 ms vs 105-140 ms) and reach condition (*stimL reachL, stimR reachR, stimL reachR and stimR reachL*). Follow-up comparisons (Šidák corrected) show that, when the right target was presented, EMG in the 105-140 ms window was significantly larger than in the 70-105 ms window in both pro-reach and anti-reach conditions (both *P* < 0.001, **Figure 5E**). When the left target was presented, EMG in the 105-140 ms window was in general smaller than in the 70-105 ms, however this was only significant in the pro-reach condition (*P* = 0.013) and not for the anti-reach condition (*P* = 0.236, **Figure 5E**). We also observed no strong correlation between the amplitude of the EMG in these two time-windows (**Figure 5F**), which suggests that the strength one response does not influence the strength of the other response. Furthermore, we saw no significant differences in express EMG magnitude (70-105 ms) between pro- and anti-reach conditions or in reciprocal EMG (105-140 ms) between pro- and anti-reach conditions (all comparisons, *P* > 0.600, **Figure 5E**).

**Figure 5.**
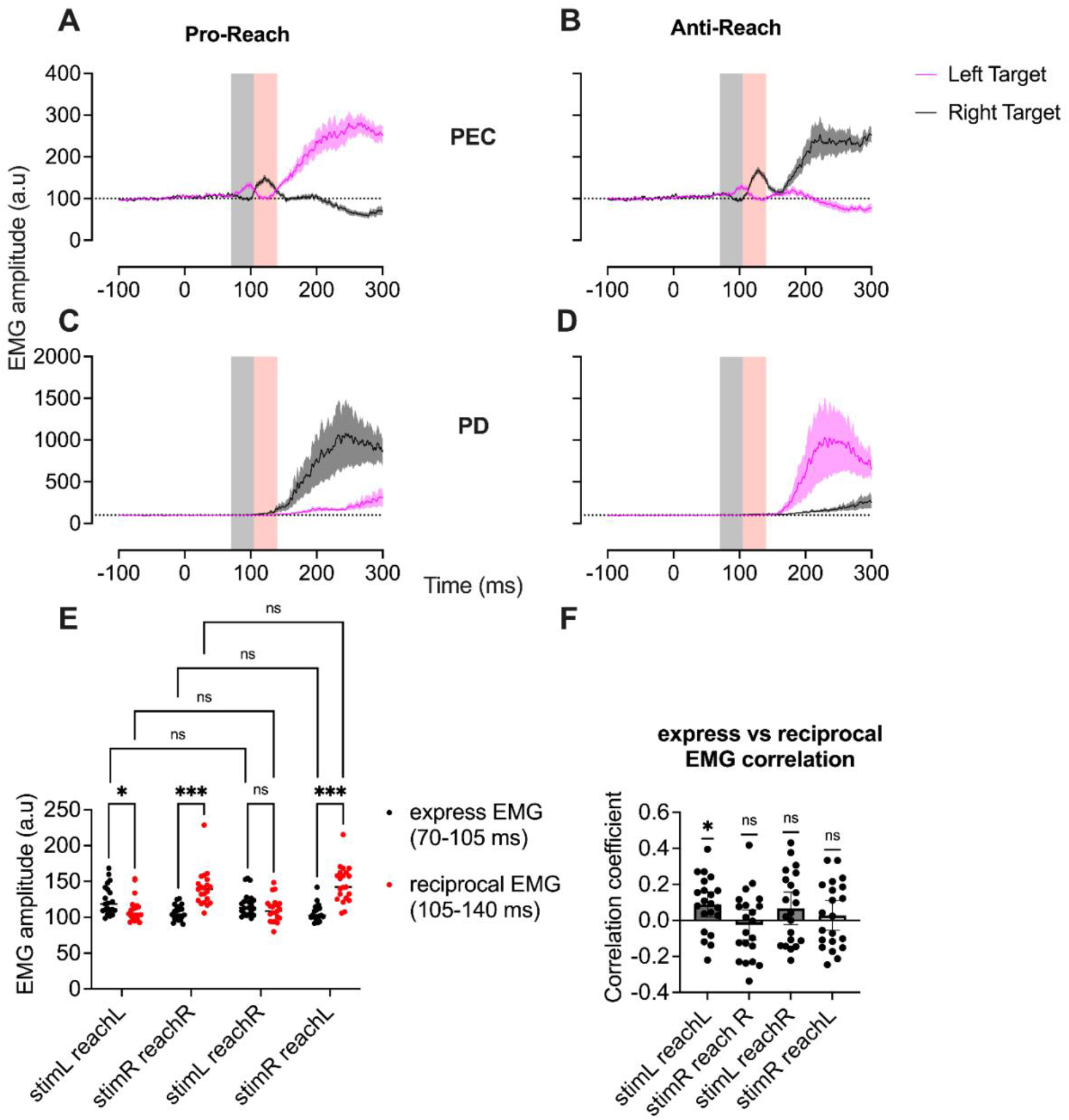
***A&B*:** Average (normalised) EMG in the pectoralis major (PEC) muscle across all subjects ***C&D*:** same in the posterior deltoid (PD) muscle. The black shaded region (70-105 ms) shows the express response window and the red shaded region shows the reciprocal response window (105-140 ms). ***E*:** Comparison between the average EMG in the express response window and the reciprocal response window. ***F*:** Correlation between the EMG amplitude in the express response window and the reciprocal response window. * P < .05, ** P < .01, *** P < .001

This confirms our qualitative impression from the exemplar raster plots that the isolated muscle activity prior to 140 ms is similar across pro-and anti-reach conditions. Note that we did not observe any early target-related responses in the posterior deltoid muscle (**Figure 5C, D**). This is not surprising as, unlike the pectoralis muscle, the posterior deltoid muscle was not preloaded.

As evident from **Figure 3** and **Figure 4**, participants did not produce a stereotypical express visuomotor response in this paradigm, instead they produced biphasic or sometimes multiphasic responses (see *stimR reachR* and *stimR reachL* in **Figure 3B, D** and **Figure 4C, D, E**). Such multiphasic responses starting in the express response window have been previously reported (Wood et al., 2015). Due to the long multiphasic nature of express visuomotor responses, it is difficult to define a set time window to check for the trial-by-trial prevalence of the express visuomotor response in this paradigm. This is especially true since some participants did not show the “classic” target-oriented activity associated with the express visuomotor response and instead showed strong reciprocal muscle responses that are also “time-locked” to the stimulus onset. If we consider the entire 70-140 ms window where the biphasic responses occur, a single trial express visuomotor response detection algorithm (Contemori et al., 2021) shows that express muscle responses were present in approximately 41% of trials across all participants. If we only consider the 70-105 ms window for the first target-oriented phase, the prevalence drops to 20%. This shows that defining the express visuomotor response as a strictly target-oriented muscle response is an incomplete or inaccurate notion. The muscle activity at 105-140 ms, despite being tuned in the direction “opposite” from the target, is still temporally coupled to the target onset and thus should be considered as part of the express visuomotor response.

To summarise, withholding the information about the rule that defines the task-goal until the appearance of the target delays the mechanical reaction time and reveals a multiphasic express visuomotor response that is nearly identical in both pro- and anti-reach conditions. The first phase (70-105 ms) of the express visuomotor response is directed at the visual target but is often weaker and less consistent than the subsequent phase (105-140 ms) that is directed in the direction opposite the target. This phasic response seems to be more strongly elicited by right than left targets.

### Experiment 2

Experiment one established that express visuomotor responses can be temporally separated from late voluntary muscle responses during visually guided reaching. In experiment two, we investigated whether target-driven early excitability changes can be temporally separated from the late movement-related excitability changes. We used the same paradigm as in experiment one and recorded Motor Evoked Potentials (MEPs) from pectoralis muscles after stimulating the left motor cortex at a range of latencies: from 60 ms after stimulus onset to just before the voluntary EMG onset. MEPs evoked from an exemplar participant are shown in **Figure 6**.

**Figure 6.**
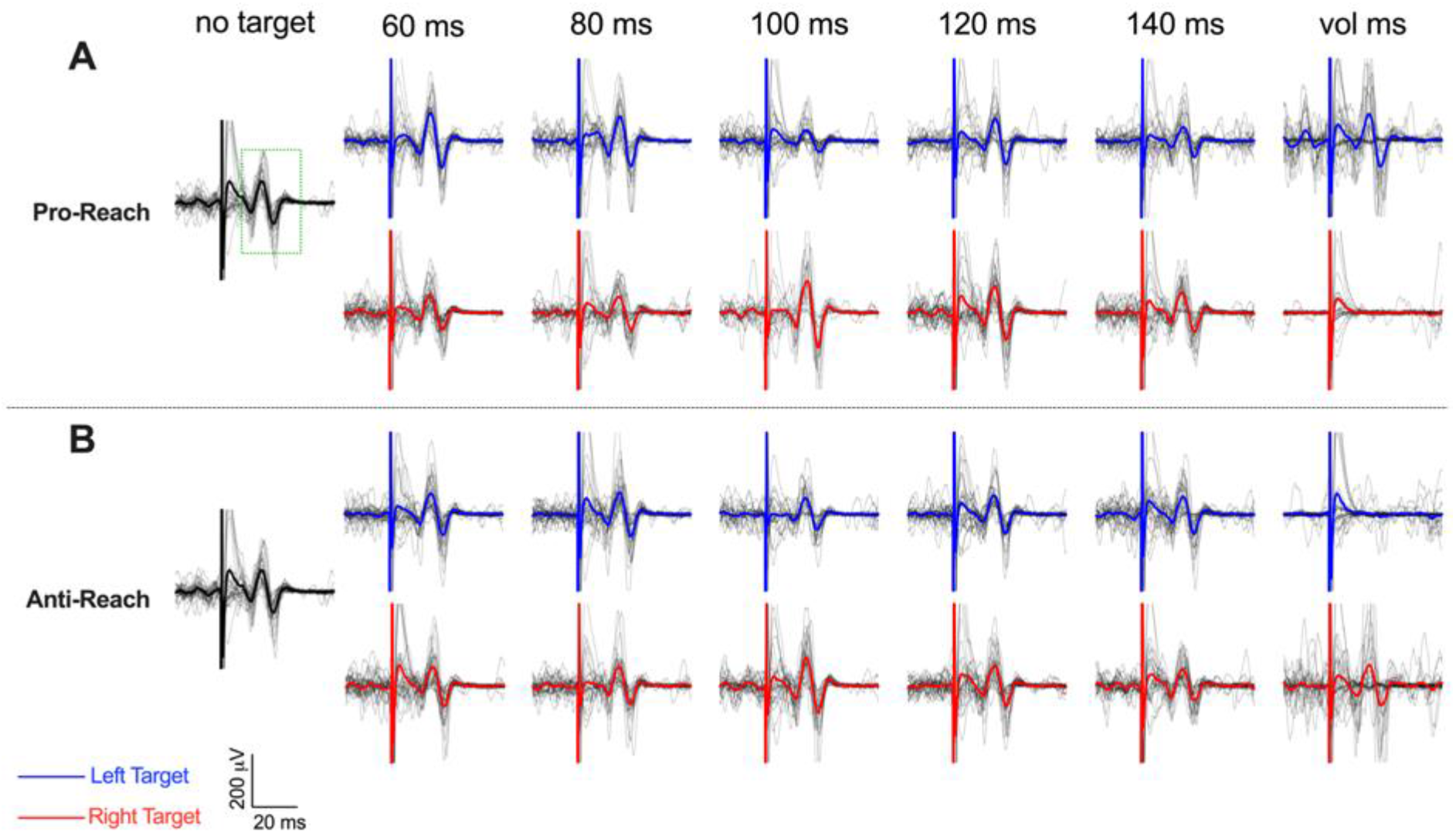
Individual MEP traces from the pectoralis major muscle of an exemplar participant with the average MEP waveform overlaid. ***A*:** The top two rows show traces from the Pro-Reach condition. ***B*:** The bottom two rows show traces from the Anti-Reach condition. The green dotted box in the no target condition for Pro-Reaches shows the time-window that was visually identified for calculating the MEP amplitude.

Like experiment one, the reaction time in experiment two was slow. The median reaction time in experiment two was ∼300 ms [range: 250–432 ms], which was not significantly different from the reaction time in experiment one (*t*_20_ = 0.623, *P* = 0.537). Introducing the magnetic stimulation did not influence the reaction time, as there was no significant difference in reaction times between stimulation (median: ∼301 ms) and non-stimulation (median: ∼304 ms) trials (*t*_20_ = 1.624, *P* = 0.120).

Due to the delayed reaction times, there was minimal movement-related background EMG for the pectoralis major muscle in the measurement windows prior to 120 ms (**Figure 7C, D**). This is in stark contrast to our previous results (cite your other paper), where substantial movement related EMG was present in the express time window (< 120 ms). The background EMG in the pectoralis muscle was influenced by the target location and the measurement window in both pro- and anti-reach conditions (Mixed-effects model; pro-reach: *F*_1.7,28.2_ = 40.2, *P* < 0.001, anti-reach: *F*_2.0,32.6_ = 50.0, *P* < 0.001). Follow up comparisons show that background EMG in the 100 ms measurement window was significantly larger for the right target than the left target in both pro- (*P* = 0.006) and anti-reach conditions (*P* = 0.003). This observation is consistent with experiment one, where we saw strong reciprocal muscle activation for the right target between 105-140 ms (**Figure 5A, B**). The background EMG measured in the late *voluntary* measurement window was consistent with task demands; compared to the right target background EMG, left target background EMG was larger in pro-reaches and smaller in anti-reaches (both comparisons *P* < 0.001).

**Figure 7.**
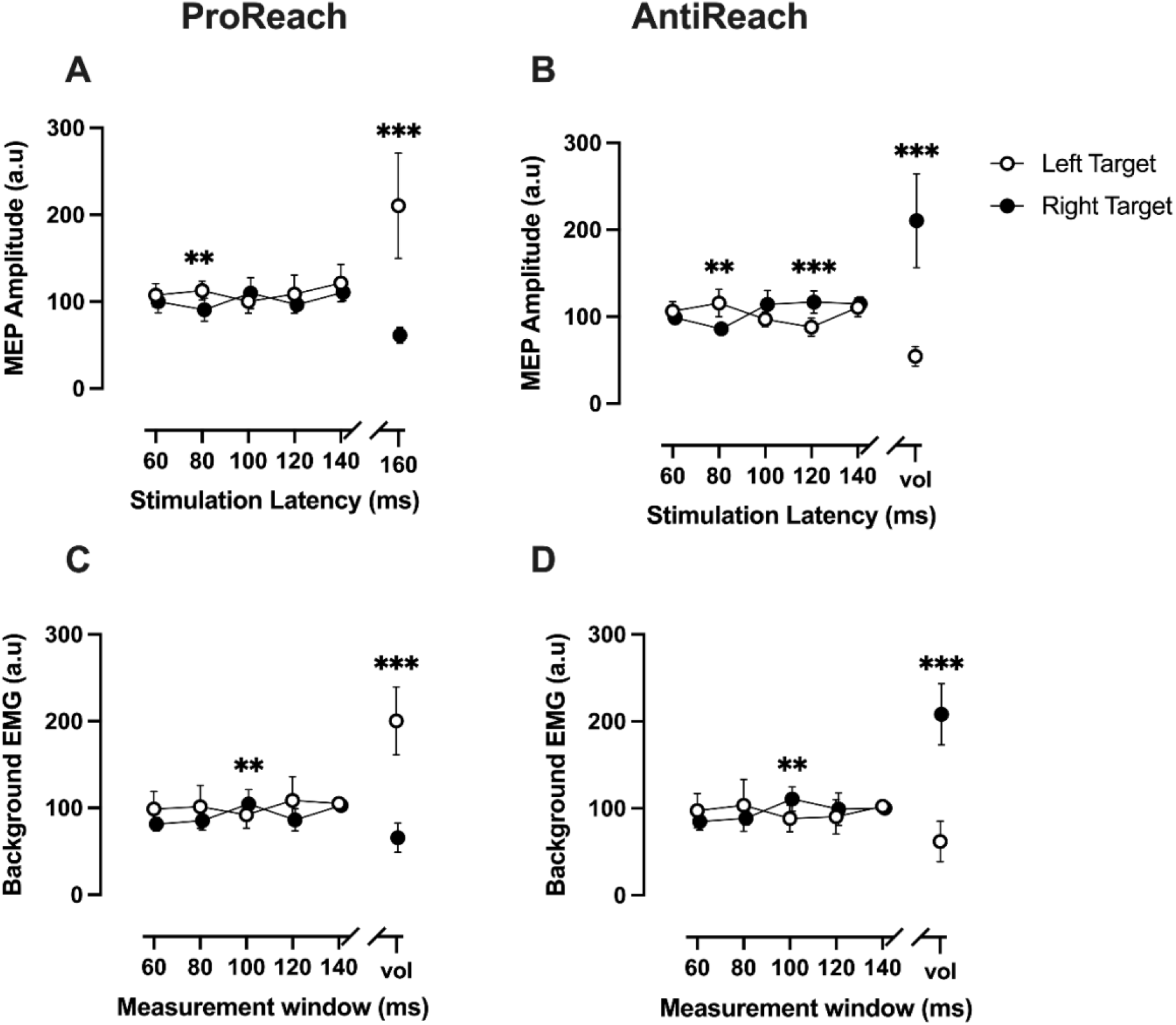
MEP and background EMG recorded from the pectoralis major muscle.***A & B*:** Time course of MEP amplitude as a function of stimulation latency. ***C & D*:** Time course of background EMG as a function of measurement window. The first column shows results in the pro-reach condition and the second column shows results in the anti-reach condition. Black circles represent the left stimulus direction and white circles represent the right stimulus direction. Error bars show 95% CI. *P < 0.05, **P < 0.01, ***P < 0.001.

In both pro- and anti-reach conditions, motor evoked potentials in the pectoralis muscle were influenced by the target location and the stimulation timing (Mixed-effects model; pro-reach: *F*_1.5,23.9_ = 20.1, *P* < 0.001, anti-reach: *F*_1.4,22.3_ = 31.3, *P* < 0.001). Follow up pairwise comparisons showed that corticospinal excitability at 80 ms was influenced by the location of the target in both pro-(*P* = 0.005, **Figure 7A**) and anti-reach conditions (*P* = 0.007, **Figure 7B**). This is consistent with the early target-oriented excitability changes observed earlier (Divakar et al., 2022), confirming our finding that the visual information has rapid access to the corticospinal tract as early as 80 ms. In that paper, we saw that the target-related MEP changes at 80 ms were immediately followed by the movement-related changes at 90-100 ms. By contrast, here we saw a significant temporal gap between the early target-related and late movement-related excitability changes. In the pro-reach condition (**Figure 7A**), after the target-oriented modulations at 80 ms, there were no significant modulations at any other early (100, 120 or 140 ms) stimulation latencies (all *P* > 0.420). When stimulation was closer to the onset of the reach, however, (at the *voluntary* latency) MEPs were significantly biased in the direction of the reach goal (*P* < 0.001). This illustrates a significant temporal gap between the early target-driven signal and the subsequent larger movement-related signal in the pro-reach condition.

The early MEP responses in the anti-reach condition were slightly different to those in the pro-reach condition (**Figure 7B**). Like the pro-reach condition, there was target-biased modulation at 80 ms. Unlike the pro-reach condition, however, there was also significant modulation at 120 ms. The right target MEPs were significantly larger than the left target MEPs at 120 ms (*P* < 0.001). It is surprising that this difference at 120 ms only occured in the anti-reach condition and not in the pro-reach condition, as there was no significant difference in the arm reaction time between pro-(∼301 ms) and anti-reach conditions (∼304 ms) (*t*_20_ = 1.609, *P* = 0.123). However, if we look at the background EMG in the anti-reach trials (**Figure 7D**), the EMG at 100 and 120 ms was generally larger for the right target compared to the left target. Thus, the pattern of excitability seems to reflect the background EMG even if the EMG effects at this latency weren’t significantly different. Interestingly, the MEP divergence at 120 ms did not continue, and disappeared by 140 ms (*P* = 0.918). The return to baseline levels at 140 ms shows that the apparently “goal-directed” modulation at 120 ms was not due to the arrival of movement-related cortical volleys, as the cortical drive prior to movement would be expected to increase over time and cause the MEP curve to diverge even more. Instead, it likely represents a part of an early alternating wave of activity evoked by target onset, like that seen in experiment one.

As is clear in **Figure 7**, the MEP and background EMG in this experiment show an overall alternating pattern similar to that in experiment one (**Figure 5A, B**). While the effect is clearer in the anti-reach condition, the general pattern of alternation is also visible in the pro-reach condition. This overall similarity in the patterns of MEP and background EMG in the early stages of reaching shows that, in this paradigm, the state of the corticospinal tract in the early stages of reaching was only influenced by express target-related signals and not by goal-directed movement signals.

## DISCUSSION

The aim of this study was to characterise the express visuomotor responses and their associated corticospinal excitability changes in isolation from subsequent voluntary muscle activity. This was achieved using a modified pro-anti reach paradigm, where the target luminance (dark or bright) conveyed the task information (pro-or anti-reach). The paradigm effectively delayed the voluntary reach and confirmed that the early target-driven responses are temporally separable from the late movement-related responses. The data show that the express and voluntary responses are indeed distinct signals, supporting the hypothesis that at least two separate neural pathways (one slow and one fast) are involved in visually guided reaching.

The modified pro-anti reach paradigm used in this study successfully delayed the voluntary reach initiation time by approximately 90 ms. In the first experiment, when the voluntary movement was delayed, express visuomotor responses appeared as a series of muscle activations and muscle inhibitions (**Figure 3, Figure 4**). The first phase of this oscillatory response typically occurred in the 70-105 ms time window and was tuned in the direction of the visual target, while the second phase typically occurred in the 105-140 ms window and was tuned in the direction opposite of the target. The phases that came after this continued in an alternating pattern until the voluntary EMG arrived. Such alternating express muscle responses were reported previously by Wood et al. (2015). Those reported express visuomotor responses that started around 70-80 ms, propagated at a frequency of 10-15 Hz, and lasted over 200 ms post stimulus onset. Since then, Atsma et al. (2018) and Kozak et al. (2019) have also reported oscillating muscle responses that started in the express response window. In these studies, the authors reported consistent target-oriented responses before the second “rebound” or “reciprocal” phase occurred. They also found the reciprocal phase to be more pronounced than the first response. In line with this, we also found the “rebound” or “reciprocal” activity to be more prominent than the preceding target-oriented response. However, contrary to the previous findings, we saw that the reciprocal EMG activity can occur even in the absence of the first target-oriented phase. This is surprising if we consider the second phase to be a consequence of a “rebound” effect from the initial wave of excitation or suppression. If that were true, the second phase should not exist without the initial response and its magnitude should be strongly correlated to the size of the first response. However, we found neither to be true. Not only did the first and second response phases appear to be causally independent (**Figure 4**), their amplitudes were unrelated (**Figure 5F**).

In experiment two, neither the MEP nor the background EMG was influenced by goal-directed activity in the early stages of reaching (**Figure 7**). This confirms that the early excitability changes that we reported previously (Divakar et al., 2022) were due at least in part to the express visuomotor response and not due solely to an overlap with the early voluntary response. The results from the two experiments in the current study were also consistent, in that both MEPs and background EMG prior to 140 ms appeared to show an alternating pattern in both pro- and anti-reach conditions, with clear target-oriented MEP modulation at 80 ms (**Figure 7**). After the initial target-driven modulation, however, there was no task specific modulation of the corticospinal tract until after 140 ms, demonstrating that the corticospinal tract receives two temporally distinct motor signals during this reaching paradigm. An express signal that generates a short-latency target directed response is followed by alternating sequences of muscle activity, and ultimately a later signal that ultimately drives a strategic goal-directed muscle response. Furthermore, in both experiments, the behaviour of the motor system in the early stages of reaching was similar across the pro- and anti-reach conditions (**Figure 5, Figure 7**), which is in stark contrast to the task-specific motor behaviour around the late voluntary movement stage. This suggests that, unlike the late response, the express oscillatory responses do not serve a direct functional purpose, and likely reflect “low-level” dynamics intrinsic to the circuits that generate express behaviour.

We and others have proposed that the express muscle response is produced by the tecto-reticulo-spinal pathway, while the late muscle response is produced by a transcortical pathway (Gu et al., 2016; Glover and Baker, 2019; Contemori et al., 2022). The rapid onset and target-oriented nature of the express responses seen in this study are consistent with a direct visuomotor transformation via the short tecto-reticulo-spinal pathway, while the more variable onset time of the late muscle response is consistent with the time delays associated with cortical processing. The origin of the oscillatory nature of express responses seen in this study is not clear, but could be a consequence of reciprocal inhibitory networks within the superior colliculus (Munoz and Istvan, 1998; Phongphanphanee et al., 2014). It seems possible that a reciprocally inhibiting mechanism, similar to the half-centre model of central pattern generation (Marder and Calabrese, 1996; Daun et al., 2009; Kohler et al., 2020), could underlie the oscillatory muscle response. According to half-centre model, when two modules of neurons (say *A* and *B*) which are under sustained excitatory drive mutually inhibit each other, then an increase in activity of one module (say *A*) would strongly inhibit the activity of the other (*B*). Subsequent self-inhibition mechanisms reduce the activity of *A* (Kohler et al., 2020), which causes a “rebound” excitation of *B* stemming from the release of the reciprocal inhibition. This leads to an alternating pattern of excitation and inhibition between the two modules of neurons. All of the necessary components to produce alternating activity by this type of mechanism are present in the superior colliculus. Studies in nonhuman primates have shown that, when a visual stimulus can appear at two potential locations in the visual field, activity in the superior colliculus map that correspond to these locations builds up prior to stimulus onset (Trappenberg et al., 2001; White and Munoz, 2011). This build-up could potentially act as the source of tonic excitatory drive required by the model. Furthermore, long range inhibitory connections are known to exist within superior colliculus (or tectum), both within and across the two tectal hemispheres (Mice: (Sooksawate et al., 2011; Phongphanphanee et al., 2014; Essig et al., 2021); Cats: (Appell and Behan, 1990; Olivier et al., 2000; Takahashi et al., 2010); Ferret: (Meredith and Ramoa, 1998; Behan et al., 2002); Monkeys: (Munoz and Istvan, 1998)). Hence, we propose a possible mechanism where, prior to target presentation, there is build-up of activity in two-mutually inhibiting the collicular regions. Presentation of the visual target triggers a series of reciprocal inhibition and rebound excitations that gets translated by the brain stem into oscillatory express visuomotor response seen in this study. Clearly, the superior colliculus is only one possible site at which alternating activity could be generated, and it remains possible that other neural substrates such as the reticular formation or spinal cord are involved. We think that a spinal origin is less likely than a midbrain or brainstem origin, however, because the second phase of the response is not dependent on the presence of overt muscle activity in the first phase. This suggests an alternation at some distance from motoneuronal activation, consistent with premotor circuits in the brain rather than the spinal cord.

## CONCLUSION

This study confirms that the earliest modulation of target-oriented corticospinal excitability during reaching is produced by circuits that underlie the express visuomotor response and not by circuits that drive fast voluntary responses. It also shows that, in a visually guided reaching task, the onset of a visual stimulus gives rise to several temporally discreet motor signals. The early alternating signals are tuned by the direction of the visual stimulus and are temporally distinct from a late muscle response that is tuned to the direction of the motor goal. The temporal gap between the express responses and the late responses suggests the involvement of two distinct (fast and a slow) neural pathways in producing these motor outputs. The properties of the fast pathway are consistent with a tecto-reticulo-spinal pathway, while those of the slow pathway are consistent with a transcortical loop.

## Notes

### Competing Interest Statement

The authors have declared no competing interest.

